# Nipah virus Bangladesh infection elicits organ-specific innate and inflammatory responses in the marmoset model

**DOI:** 10.1101/2021.10.11.463955

**Authors:** Christian S. Stevens, Jake Lowry, Terry Juelich, Colm Atkins, Kendra Johnson, Jennifer K. Smith, Maryline Panis, Tetsuro Ikegami, Benjamin tenOever, Alexander N. Freiberg, Benhur Lee

## Abstract

The common marmoset (*Callithrix jacchus*) is increasingly recognized as an ideal non-human primate (NHP) at high-biocontainment due to its smaller size and relative ease of handling. Here, we evaluated the susceptibility and pathogenesis of Nipah virus Bangladesh strain (NiV_B_) infection in marmosets at biosafety level 4. Infection via the intranasal and intratracheal route resulted in fatal disease in all four infected marmosets. Three developed pulmonary edema and hemorrhage as well as multi-focal hemorrhagic lymphadenopathy, while one recapitulated neurologic clinical symptoms and cardiomyopathy on gross pathology. Organ-specific innate and inflammatory responses were characterized by RNA-seq in six different tissues from infected and control marmosets. Notably, a unique transcriptome was revealed in the brainstem of the marmoset exhibiting neurological symptoms. Our results provide a more comprehensive understanding of NiV pathogenesis in an accessible and novel NHP model, closely reflecting clinical disease as observed in NiV patients.

## Background

Nipah virus (NiV), a zoonotic, bat-borne negative-sense paramyxovirus of the genus *Henipavirus*, has seen outbreaks in humans nearly every year since 1998. There have been over 600 cases of respiratory and/or neurological disease in patients, 59% percent of which resulted in death [1–3]. All but two major outbreaks have been caused by NiV-Bangladesh (NiV_B_), which has had a higher case fatality rate (CFR) of 76% relative to the other NiV clade, NiV-Malaysia (NiV_M_), which has had a CFR of 39% [2]. NiV_B_ has a shorter average incubation period, higher rate of respiratory symptoms, and smaller proportion of patients exhibiting segmental myoclonus relative to NiV_M_ [4–6]. Human-to-human transmission has also been seen more commonly in NiV_B_ outbreaks but extremely rare in NiV_M_ outbreaks [7]. Given these significant differences animal models successfully recapitulating NiV_B_ pathology in humans are especially critical.

Several non-human primate (NHP) models have been used to investigate NiV pathogenesis and vaccine research: African green monkeys (*Chlorocebus aethiops;* AGM), cynomolgus macaques (*Macaca fascicularis*), squirrel monkeys (*Saimiri sciureus*), and grivets (*Chlorocebus aethiops*) [8,9,18–22,10–17]. Among these, only AGMs faithfully recapitulate the high lethality of NiV_B_ exposure as well as most clinical signs observed in patients. However, overt clinical signs of neurological disease are seen very rarely in AGMs despite neurological disease being commonly found via MRI and post-mortem histopathology. Additionally, myocarditis has been consistently reported among patients infected with NiV_B_ but reports of myocarditis in animal models are rare [23].

AGMs are still able to replicate much of the other hallmark clinical pathology and histopathology of NiV infection: systemic vasculitis, hypoalbuminemia, pneumonitis, coagulopathy, and congestion in the liver and spleen [9,18,21]. Transcriptional profiling in AGMs has also shown significant upregulation in innate immune signaling genes (*MX2, OASL, OAS2*), particularly among non-survivors of infection relative to survivors [21]. Others have shown that NHP models show an especially robust expression of interferon stimulated genes in the lung, including *ISG15* and *OAS1* [24].

There has been notable success using AGMs as a model for NiV_B_ infection, especially in the evaluation of vaccine candidates and therapeutics [19,22]. However, the lack of clinical neurological symptoms and myocarditis, the logistical difficulties associated with AGMs’ larger size, and the shortage of animals due to COVID19 research, call for a diversification in the preclinical testing portfolio of NHP models. We show here that NiV_B_ infection of the common marmoset (*Callithrix jacchus*) can faithfully recapitulate pathology seen in humans. Marmosets were chosen due to several distinct advantages. First, they are an ideal NHP model for studying high-biocontainment human respiratory pathogens given their small size and relative ease of handling. Second, they are the first among New World monkeys to have their whole genome sequenced and assembled allowing for RNA-sequencing (RNA-seq) to elucidate transcriptomic changes in infected and uninfected tissues [25]. Lastly, transmission experiments are made easier given the requirement of co-housing, and their high reproductive efficiency and short gestation period and delivery interval allows for easier development of transgenic animals [26]. With these advantages in mind, results of this study identified marmosets as a highly susceptible novel New World NHP model of NiV_B_ infection and pathogenesis.

## Methods

### Ethics statement

Experiments were approved by the Institutional Animal Care and Use Committee of the University of Texas Medical Branch (UTMB) and performed following the guidelines of the Association for Assessment and Accreditation of Laboratory Animal Care International (AAALAC) by certified staff in an AAALAC-approved facility. Animals were co-housed allowing social interactions, under controlled conditions of humidity, temperature and light. Animals were monitored pre- and post-infection and fed commercial monkey chow, treats and fruit twice daily. Food and water were available ad libitum and environmental enrichment consisted of commercial toys. Procedures were conducted by BSL4-trained personnel under the oversight of an attending BSL4-trained veterinarian. All invasive procedures were performed on animals under isoflurane anesthesia. Animals that reached euthanasia criteria were euthanized using a pentobarbital-based solution.

### Virus and cells

NiV_B_ (200401066 isolate) was kindly provided by Dr. Thomas Ksiazek (UTMB). The virus was propagated on Vero E6 cells (ATCC, CRL1586) and virus titers were determined by plaque assay on Vero cells (ATCC, CCL-81) as previously described [27]. All work with infectious virus was performed at BSL4 biocontainment at the Galveston National Laboratory at UTMB.

### Animal studies

Four healthy marmosets (2 females, 2 males; 348-453 grams) were inoculated with 6.33E+04 plaque forming units (PFU) of NiV_B_, dividing the dose equally between the intranasal (IN) and intratracheal (IT) routes for each animal. After infection, animals were closely monitored for signs of clinical illness, changes in body temperature (BMDS transponders) and body weight. A scoring system was utilized monitoring respiratory, food intake, responsiveness, and neurological parameters to determine study endpoint. In addition, changes in blood chemistry and hematology were monitored. Blood was collected via the femoral vein.

### Hematology and serum biochemistry

Blood was collected in tubes containing EDTA and complete blood counts (CBC) were evaluated with a VetScan HM5 (Abaxis). Hematological analysis included total white blood cell counts, white blood cell differentials, red blood cell counts, platelet counts, hematocrit, total hemoglobin concentrations, mean cell volumes, mean corpuscular volumes and mean corpuscular hemoglobin concentrations. Clinical chemistry analysis was performed using a VetScan2 Chemistry Analyzer. Serum samples were tested for concentrations of albumin (ALB), amylase (AMY), alanine aminotransferase (ALT), alkaline phosphatase (ALP), glucose (GLU), total protein (TP), total bilirubin (TBIL), plasma electrolytes (calcium, phosphorus, sodium, and potassium, blood urea nitrogen (BUN), creatinine (CRE), and globulin (GLOB)).

### X-ray images

X-ray images were obtained using the MinXray model HF 100/30 portable x-ray unit (MinXray) operated at 78 kVp 12 3.20 mAs, 40” from the subject. A Fuji DR 14×17” panel served as the exposure plate and the images were processed in the SoundVet application (SoundVet).

### Sample collection and RNA isolation

Whole blood was collected in EDTA Vacutainers (Beckman Dickinson) for hematology and serum biochemistry, while another part was mixed with TRIzol LS for RNA extraction. At euthanasia, full necropsies were performed and tissues were homogenized in TRIzol reagent (Qiagen) using Qiagen TissueLyser and stainless-steel beads. The liquid phase was stored at −80°C. All samples were inactivated in TRIzol LS prior to removal from the BSL4 laboratory. Subsequently, RNA was isolated from blood using the QIAamp viral RNA kit (Qiagen) and from tissues using the RNeasy minikit (Qiagen) according to the manufacturer’s instructions.

### RT-qPCR

RNA was extracted using Direct-zol RNA Miniprep kits (Zymo Research). RT-qPCR was run using QuantiFast RT-PCR mix (Qiagen), probes targeting NiV_M_ P gene (5’-ACATACAACTGGACCCARTGGTT-3’ and 5’-CACCCTCTCTCAGGGCTTGA-3’) (IDT), and fluorescent probe (5’-6FAM-ACAGACGTTGTATA+C+CAT+G-TMR) (TIB MOLBIOL) RT-qPCR was performed using the following cycle: 10 minutes at 50□°C, 5 minutes at 95□°C, and 40 cycles of 10□seconds at 95□°C and 30 seconds at 60□°C using a BioRad CFX96 real time system. Assays were run in parallel with uninfected hamster tissues and a NiV_M_ stock standard curve.

### Virus titration

Virus titration was performed by plaque assay on Vero cells as previously described [27].

### Histopathology and immunohistochemistry

Tissues were immersion-fixed in 10% neutral buffered formalin for at least 21 days under BSL4 conditions with regular formalin changes following approved protocols. Specimens were then transferred to and processed under BSL2 conditions. Briefly, tissues were dehydrated through a series of graded ethanol baths, infiltrated by and embedded with paraffin wax, sectioned at 5 μm thickness, and stained with hematoxylin and eosin. Immunohistochemistry for NiV nucleoprotein detection was performed using a rabbit anti-NiV-nucleoprotein antibody incubated overnight (kindly provided by Dr. Christopher Broder, Uniformed Services University, Bethesda, Maryland) and a secondary horseradish peroxidase-conjugated goat antirabbit antibody (Abcam) incubated for 2 hours (both 1:1000). DAB substrate (ThermoScientific) was then added for 2 minutes for chromogenic detection HRP activity.

### RNA sequencing (RNA-seq)

Libraries were prepared using the TruSeq RNA Library Prep Kit v2 (Illumina) or TruSeq Stranded mRNA Library Prep Kit (Illumina). Libraries were sequenced on an Illumina NextSeq 500 platform using a paired end 2×150-base pair, dual index format. Across all marmosets five tissue samples were sequenced (lung, inguinal lymph node, kidney, brainstem, and spinal cord) with gonads sequenced only in the infected marmosets. Tissues from the naïve marmoset were sequenced in technical duplicate. Tissues from naïve marmoset were kindly provided by Dr. Ricardo Carrion Jr (Texas Biomedical Research Institute, San Antonio, Texas).

### RNA-seq data processing

Trimming, alignment, and quantification were all performed in Partek Flow v10.0 [28]. Ends were trimmed based on quality score then aligned to the *Callithrix jacchus* genome (Ensemble release 91) [29] using Bowtie2 v2.2.5. Counts were generated with *Callithrix jacchus* annotation v105 (Partek E/M). Normalization was performed by trimmed mean of M-values. Six samples were removed from the final analysis based on either having fewer than 1E+06 mapped reads or were determined to be outliers based on principal component analysis (PCA). Differential gene expression analysis was performed using Partek’s gene specific analysis (GSA) model.

Unaligned reads were then aligned to NiV_B_ (GenBank: AY988601.1) using BWA-MEM (BWA v0.7.17). Transcriptional gradient analyses was performed using IGVtools on samples with >100 aligned reads. P editing was measured using mPileup in Samtools v1.8 [30], selecting reads with inserts of Gs only and samples with >100 reads at the P editing site.

### Statistical analysis, gene set analysis, and visualization

PCA and hierarchical clustering was performed using Partek Flow v10.0. Functional enrichment analysis was performed using PantherDB’s Biological Processes Gene Ontology (GO) annotation set [31] unless otherwise noted. Gene lists were acquired via Partek’s GSA model, filtering by FDR step-up < 0.05 and fold change > 2. Gene lists were analyzed via the PANTHER Overrepresentation Test, annotation version release data 2021-08-18. Due to a lack of gene ontology sets for *Callithrix jacchus*, the *Homo sapiens* GO biological process complete annotation data set was utilized. Only GO terms where FDR < 0.01 were used. GO terms along with accompanying FDR were clustered and visualized using REVIGO [32].

## Results

### NiV_B_ infection and pathogenesis in marmosets

In the studies described here, we evaluated the susceptibility and suitability of the common marmoset to infection with NiV_B_. Four marmosets were infected with NiV_B_ by the IN/IT routes and throughout the course of the study, survival, body weights, body temperature, blood chemistry and complete blood count (CBC) were determined (**Supplementary Figures S1 and S3-S6**). All animals developed clinical signs of disease and reached euthanasia criteria between days 8 and 11 post infection (**Table 1**). The first signs of infection were seen by day 8 when subjects #300 and #425 displayed anorexia and hyperventilation. Subject #425 then became lethargic and reached euthanasia criteria. Subject #300 deteriorated on day 9, when found with open mouth breathing and reached euthanasia criteria. Subject #193 had a reduced food consumption by day 9, and by day 11 was hyperventilating and lethargic and reached euthanasia criteria. Subject #340 was normal through day 10 PI, then developed a hunched posture, and on day 11 PI was hyperventilating and reached euthanasia criteria. Furthermore, #340 presented left hindlimb tremors during euthanasia, reflecting signs of neurological impairment.

**Table 1.**
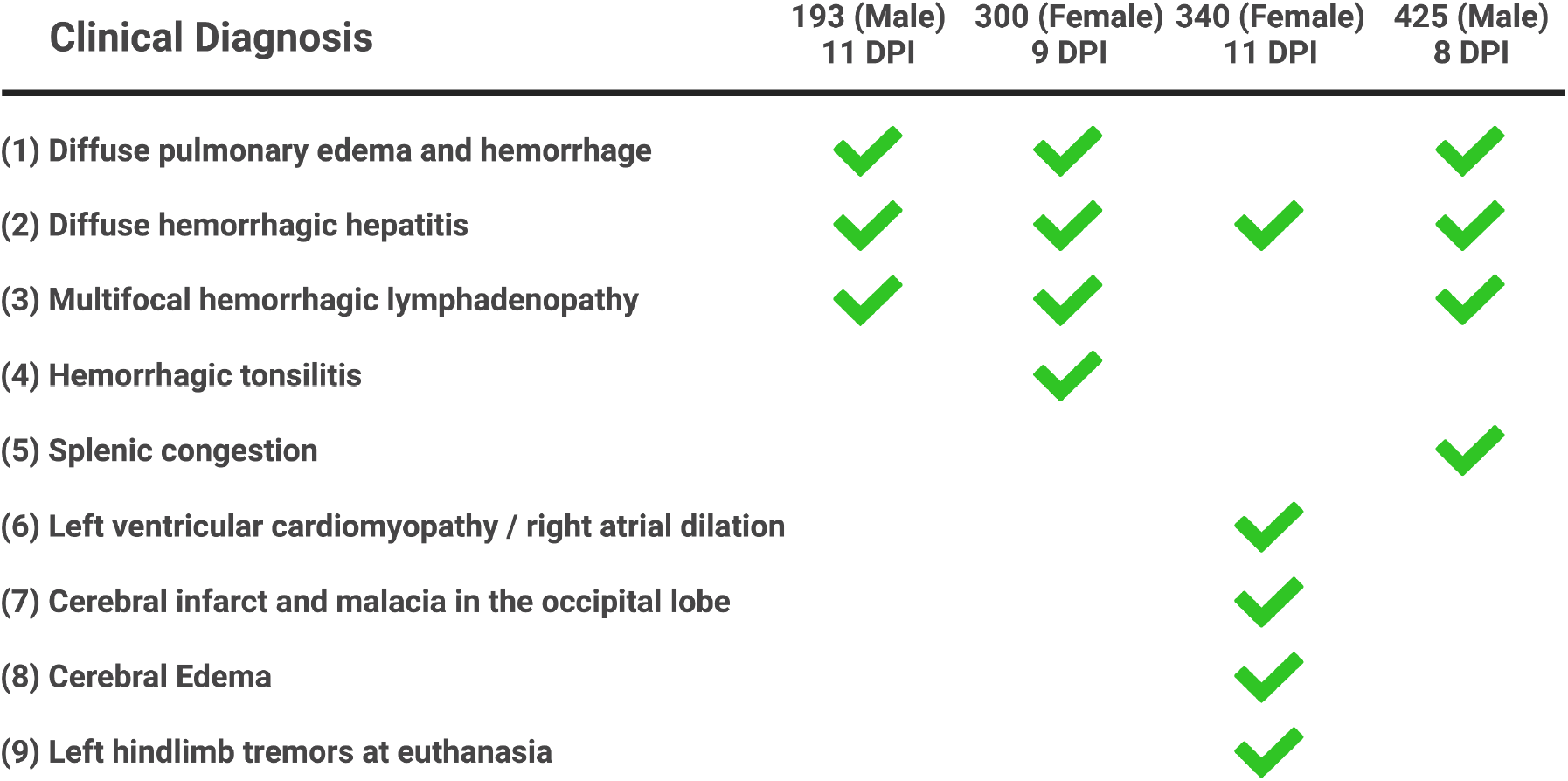
Clinical description and diagnosis in NiV_B_ infected marmosets. Graphical representation of clinical descriptions and diagnoses of all four marmosets infected with NiV_B_. Date of necropsy noted for each marmoset. Further detail is available in **Table S1**.

To monitor development of respiratory disease, we used X-ray radiographic imaging (**Figure 1A and Supplementary Figure S2**). Prior to day 7, no radiographic changes were observed in the lungs. By euthanasia, virtually no normal lung was observed for animals #425, #300, and #193. Animal #340 exhibited some inflammation around the smaller bronchi. Gross pathology found <5-10% normal lung tissues in the cranial portion of the lung (**Figure 1B and Table S1**) while the rest was filled with hemorrhagic/serous fluid. The livers had multi-focal to coalescing regions of pale circles and were very friable (**Figure 1C**). Lymph nodes had gross enlargement and hemorrhage and in animal #425, the spleen was friable and congested. Interestingly, animal #340 presented with neurological involvement as well as a less severe infection of the lungs. An infarct was found, as well as malacia near the transition zone between grey and white matter in the occipital lobe of the left cerebral hemisphere, and edema was present in the cerebrum. According to these observations, marmosets developed diffuse pulmonary edema and hemorrhage, diffuse hemorrhagic hepatitis, and multi-focal hemorrhagic lymphadenopathy. Of note, one animal (#340) had both left ventricular cardiomyopathy and neurological complications.

**Figure 1:**
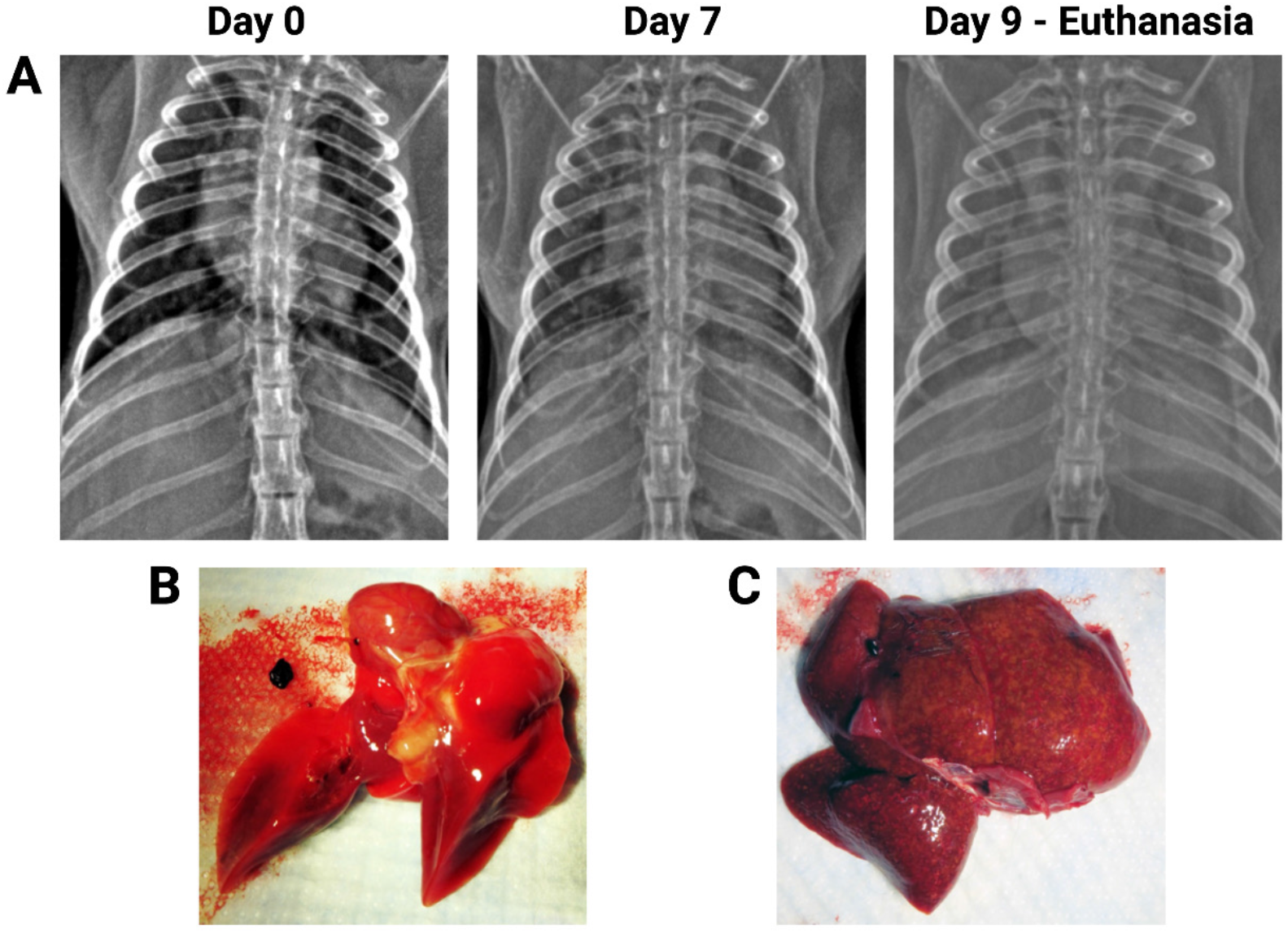
Progressive pathogenesis in marmosets after NiV_B_ infection. **(A)** Subject #300 was positioned in a ventral dorsal position and all images are in a R -> L orientation. X-ray images of the chest were taken on days 0, 1, 3, 5, 7, and at euthanasia. Prior to day 7, lungs appeared normal. Progressive lung pathology was detected from day 7 onward, with a decrease in normal lung tissue over the experimental infection course. By day 7, some decreasing opacity was observed along with an appearance of an interstitial pattern. At the time of Euthanasia (day 9 post-infection), virtually no normal lung was observed as the lungs develop a nodular alveolar pattern, suggesting the alveoli are filled with fluid. **(B)** Lung of subject #425 at time of euthanasia at day 8 post-infection. Acute diffuse pulmonary hemorrhage, with hemorrhage involving all lobes and <10% of normal tissue remaining. **(C)** Liver of subject #425. Diffuse massive necrosis of the liver, with necrosis involving all lobes and normal structure visible.

Histopathological and immunohistochemical examination of the lungs showed that all animals had mild to moderate infiltrate in pulmonary vessels (**Figure 2A**). Both syncytia formation and NiV antigen were evident in endothelial and smooth muscle cells in pulmonary vessels as well as capillary endothelial cells in the alveolar septum in all four animals (**Figure 2B**). Furthermore, histopathological lesions in subject #425 suggests myocarditis associated with the infiltration of mononuclear cells and neutrophils (**Figure 2C**). Viral antigen was found in the heart smooth muscle cells and capillary or arteriole endothelial cells (**Figure 2D**). Viral antigen was also detected in the in heart smooth muscle cells and capillary endothelial cells of #300. No detectable lesions or viral antigen were found in the testes, bronchus, or central nervous system (CNS) of any subjects. Significant histopathological changes and viral antigen were found in the liver, spleen, and kidneys of all marmosets (**Supplemental Figure S7**).

**Figure 2:**
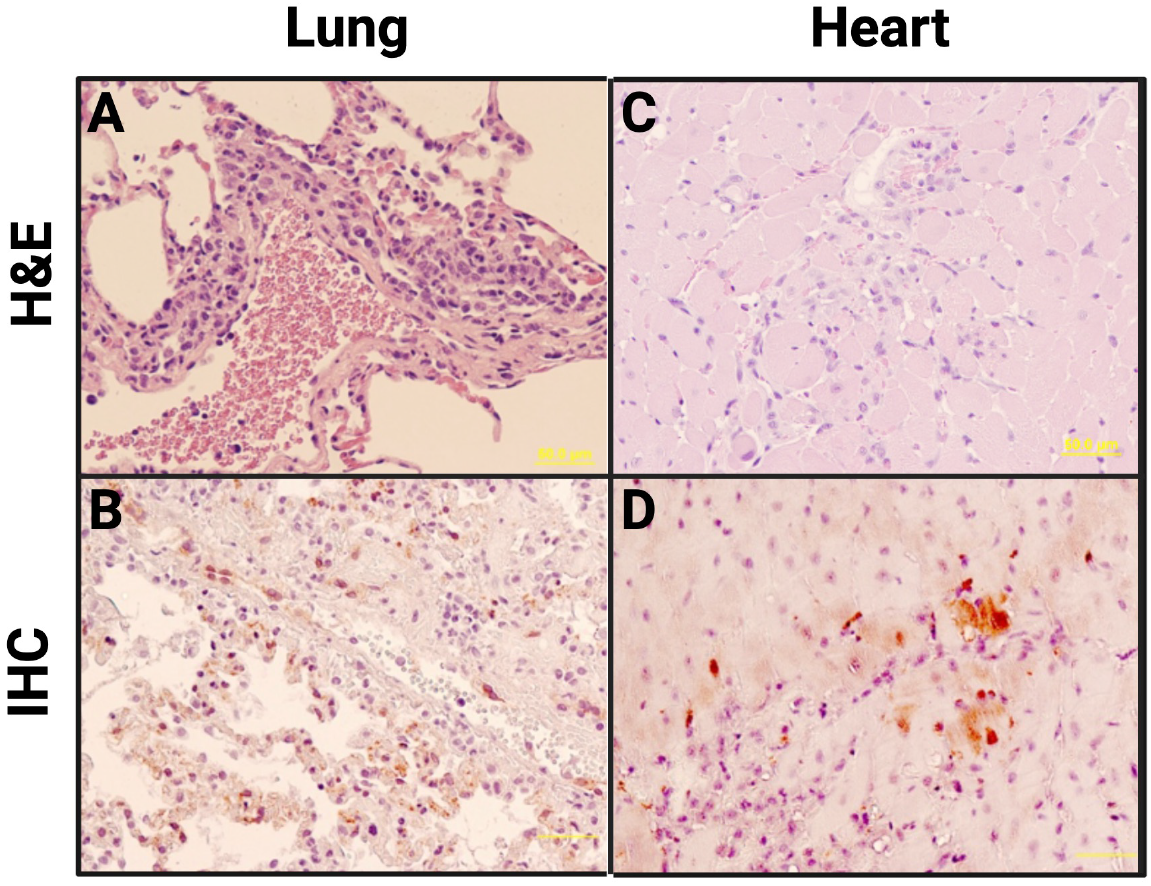
Histopathological changes in marmoset tissues after NiV_B_ infection. **Panels A and C:** H&E staining**. Panels B and D:** Immunohistochemical staining for NiV nucleoprotein. **(A)** Lung (subject #340): Mild to moderate infiltration of mononuclear cells (macrophages, monocytes) and neutrophils in perivascular space. **(B)** Lung (subject #425): NiV antigen present in endothelial cells and smooth muscle cells of pulmonary vessels (arteriole, venule). **(C)** Heart (subject #425): Mild myocarditis with necrosis of myocytes and infiltration of mononuclear cells and neutrophils. **(D)** Heart (subject #425): NiV antigen present in capillary and arteriole endothelial cells, as well as heart muscle cells.

Blood chemistry and CBC were monitored at days 1, 3, 5, 7, and 10 PI (**Supplementary Figures S3-S6**). Subject #193 showed an increased white blood cell one day before euthanasia, indicative for leukocytosis (**Supplementary Figure S3**). Neutrophilic and eosinophilic granulocytosis immediately before euthanasia was noted for subjects #193 and #300, and subjects #193, #300, and #425, respectively. Additionally, basophilia was noted for subjects #193 and #340 starting 1 DPI. Subject #425 showed marked increases in hematocrit and platelet counts one day prior to euthanasia, possibly indicating reactive thrombocytosis.

Elevations in alkaline phosphatase and/or alanine aminotransferase were suggestive of hepatocyte damage in #193, #340, and #425 (**Supplementary Figure S4**), consistent with our histopathological results and reflective of laboratory abnormalities found in NIV-infected patients. Amylase levels showed a decreased trend over the course of the study for all four subjects, indicative for damage in the pancreas (**Supplementary Figure S6**).

### Viral titers and RT-qPCR in tissues

23 tissues were collected at euthanasia in order to determine viral titer (**Figure 3A**) and measure viral RNA (**Figure 3B**). Titerable virus was found in 19 of the tissues evaluated. #425 had detectable NiV_B_ in seven of the ten tissue groups examined, #300 in six, #193 in four, and one for #340. Infectious virus was found in both the brainstem and the cerebellum of #300. Viral RNA was detected in at least one marmoset in each tissue evaluated. Viral RNA was found in the CNS for all marmosets except #193.

**Figure 3.**
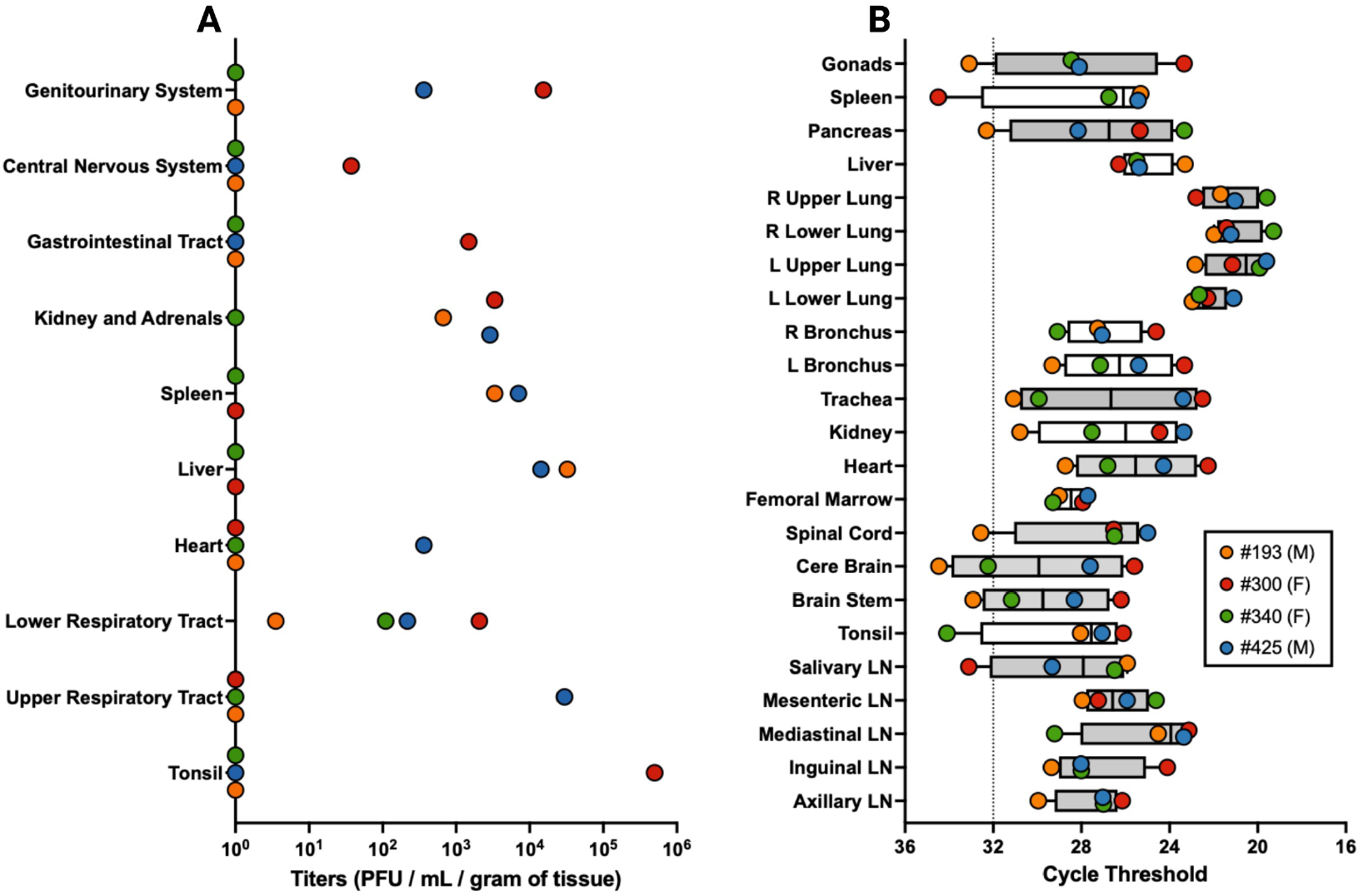
Viral titers and viral RT-qPCR data across various tissues. **(A)** At time of euthanasia, tissues were collected. Viral titers were determined by plaque assay in 23 tissues, combined into 10 tissues systems seen here. Values equal to zero were set to 10^0^ for log-scale visualization. Tissue systems with more than one tissue type are upper respiratory tract (nasal mucosa and trachea), lower respiratory tract (left upper lobe, left lower lobe, right upper lobe, and right lower lobe of the lung as well as each of the left and right bronchi), kidney and adrenals (kidney and adrenal gland), gastrointestinal tract (pancreas, jujunum, and colon transversum), central nervous system (brain-frontal, brain-cerebellum, brainstem, and cervical spinal cord), and genitourinary system (urinary bladder and gonads). **(B)** RT-qPCR was performed to determine relative amounts of viral RNA across 23 tissues. Average cycle threshold for uninfected samples is shown (dotted line). Fill color for box and whisker plots denote similar tissue types (e.g. all lymph node (LN) samples are adjacent and grey). Both right (R) and left (L) lower and upper lung samples are adjacent and also grey.

### RNA-seq of NiV_B_ and *Callithrix jacchus* in infected tissues

RNA was extracted from spinal cord, brainstem, inguinal lymph node, gonads, kidney, and lung. The average sample contained 17.4 million reads aligned to *Callithrix jacchus* and the most highly infected tissue contained >180,000 NiV_B_ reads (**Figure 4A**). In the marmoset model, NiV_B_ exhibits a transcriptional gradient that does not drop significantly until after N, P, and M (**Figure 4B**). Despite the steep transcriptional gradient after M, three samples achieved >95% coverage of L. We also measured RNA editing in P and found approximately 40% of all P-derived reads were V and W and the most Gs inserted into any read were 12 (**Figure 4C**).

**Figure 4.**
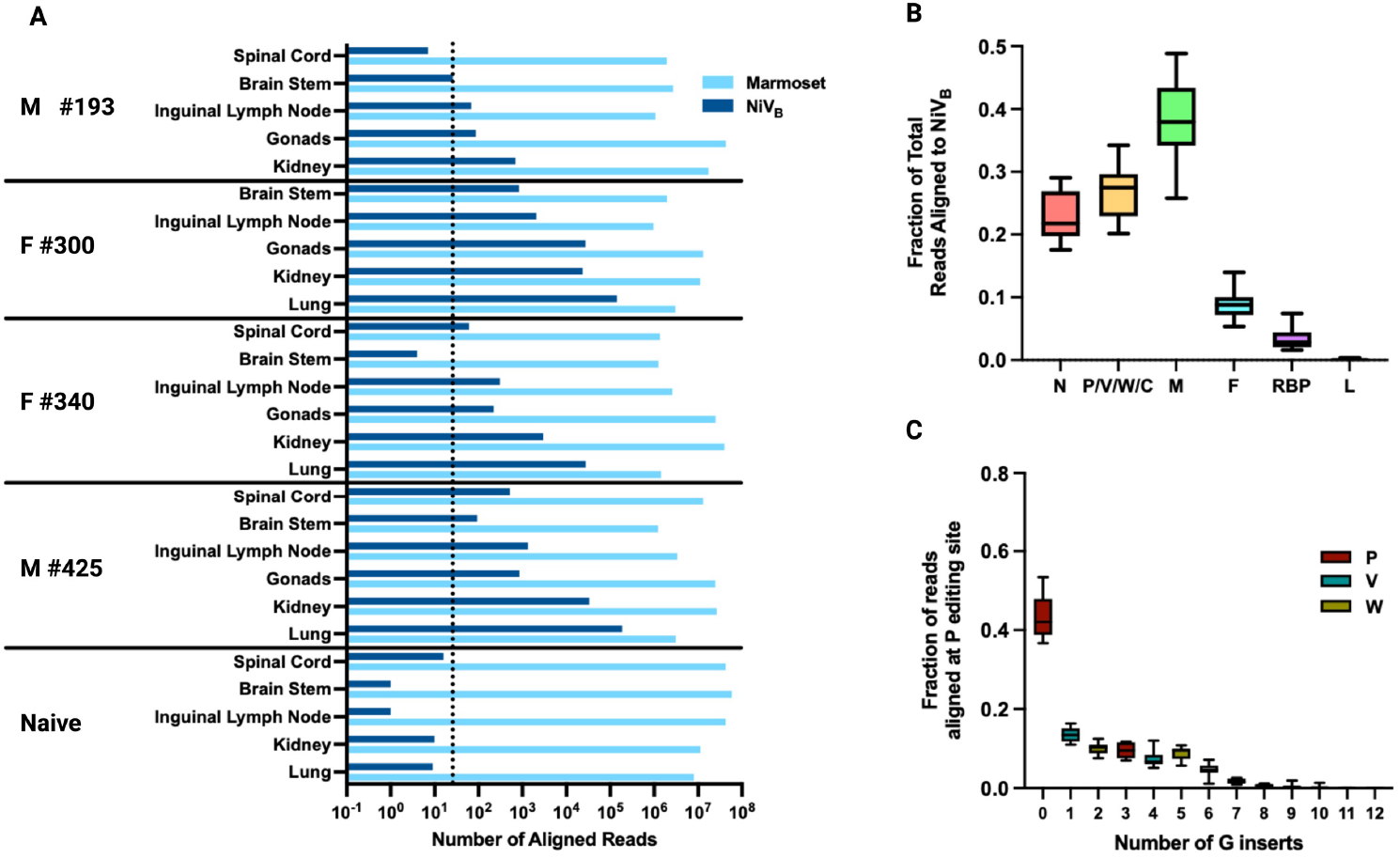
RNA-seq of NiV_B_ in infected marmoset tissues. **(A)** Total number of reads aligned to *Callithrix jacchus* (cyan) and NiV_B_ (dark blue) are shown across all tissues and samples. The dotted line is placed three standard deviations above the median number of reads aligning to NiV_B_ across all naïve samples, the threshold for inclusion into 4B. **(B)** The fraction of total reads aligning to each NiV_B_ gene across all samples exceeding 100 NiV_B_ reads. **(C)** The fraction of reads with the indicated number of inserted guanines (Gs) at the RNA editing site in P for all samples with a depth of at least 100 reads at the RNA editing site.

As expected, PCA shows that samples primarily cluster based on tissue-type (**Figure 5A**). Using differential expression analysis, 293 genes were found to be both consistently upregulated in each tissue and significantly upregulated when comparing all infected tissues against all uninfected tissues (**Figure 5B**). Enrichment analysis with these genes shows innate and inflammatory associated biological processes, in addition to processes involved in apoptosis and cell death (**Figure 5C**). Hierarchical clustering with the 25 most significant of these genes shows clustering based on both tissue-type and infection status (**Figure 5D**). Genes involved in the compliment cascade, interferon signaling, and genes in the major histocompatibility complex locus are well represented.

**Figure 5.**
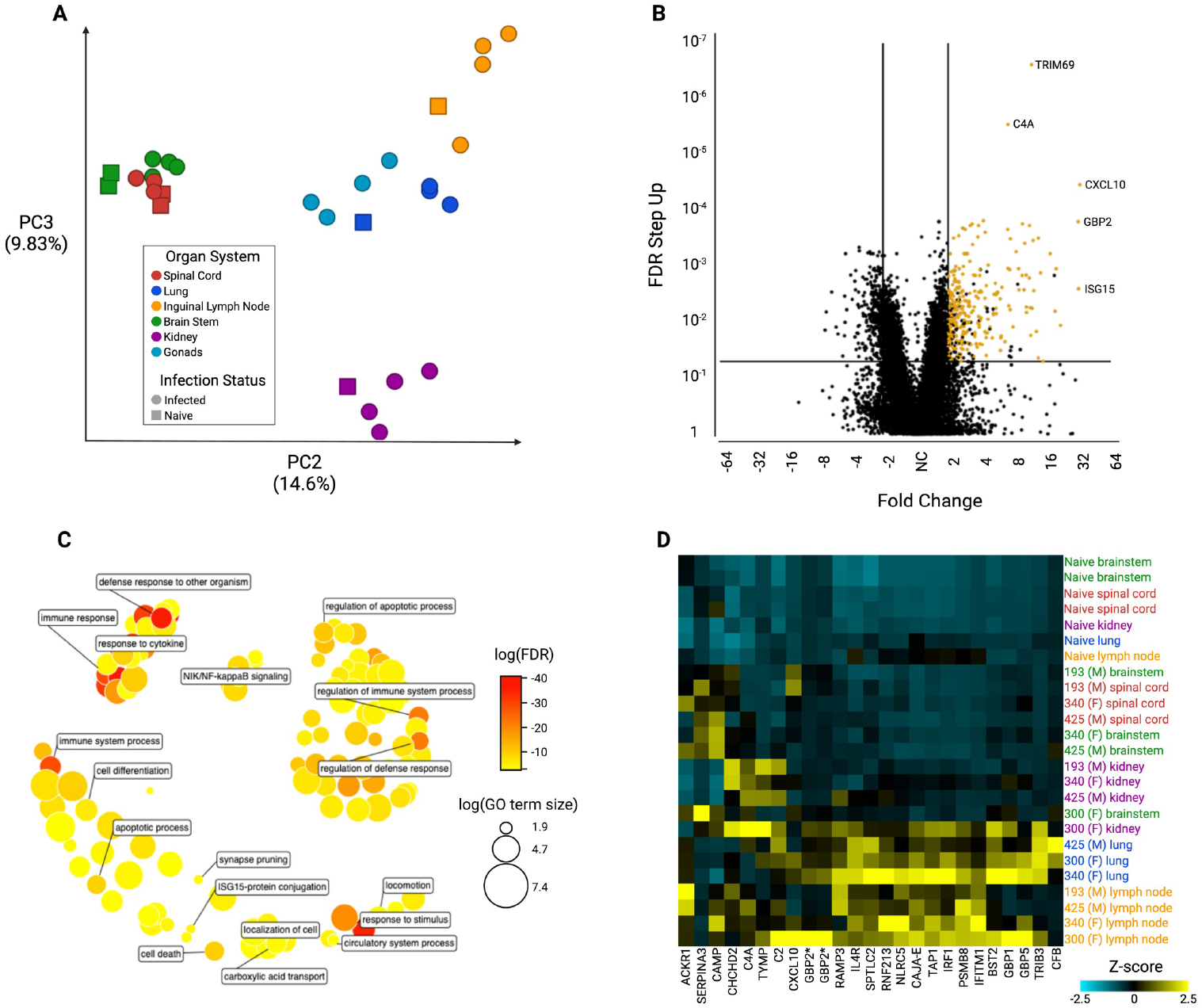
RNA-seq samples aligned to *Callithrix jacchus*. **(A)** Principal component analysis (PCA) was performed on all samples showing that organ system is the primary driver of gene expression differences between samples. Samples from similar organ systems (e.g. brainstem and spinal cord) also group together. PC1 is not shown as it is primarily driven by expression differences separating male and female gonad samples from the rest of the dataset. **(B)** A volcano plot comparing all infected samples to all uninfected samples. Lines are placed at 2 (high expression in infected samples) and −2 (higher expression in uninfected samples) as well as FDR = 0.05. The 293 genes highlighted in yellow are upregulated in each individual tissue comparison of infected versus uninfected, and in the total sample set are at least 2-fold higher expressed in infected versus uninfected with an FDR step-up value of <= 0.05. **(C)** The 293 yellow-highlighted differentially expressed genes from (B) were used to obtain enriched gene ontology (GO) gene sets, and similar terms were clustered in semantic space using REVIGO [32]. Log_10_(FDR) of the gene set is indicated by the color and log_10_ of the number of terms in the gene set is indicated by the size of the circle. **(D)** Differential expression analysis across all samples was used to identify the top 25 genes most overexpressed in infected versus uninfected samples—ranked by FDR step up value—and hierarchical clustering was performed to obtain a heatmap. Gene expression is highest in yellow samples and lowest in cyan samples. *An orthologue or “like” protein via the National Center for Biotechnology Information (NCBI).

We also focused specifically on the lungs and the brainstem. In the lungs we see interferon stimulated genes such as ISG15, OAS3, OASL, and IFNB1 upregulated in infected tissues (**Figure 6A**). Here, the high enrichment of innate and inflammatory pathways is particularly obvious (**Figure 6B**). We also found that in the brainstem of the only marmoset with neurological symptoms, #340, the transcriptome is distinct (**Figure 6C**). Using gene ontology, we found significant enrichment in terms dealing with myelination and oligodendrocyte function. We then identified OPALIN, a marker of oligodendrocytes in gene expression profiling, and OPALIN’s 15 nearest neighbors based on single cell RNA expression data according to the Human Protein Atlas [33]. Using this we show that only marmoset #340 shows high expression across all the identified genes (**Figure 6D**, magenta). These results may be noteworthy given previous findings of significant myelin degradation and axonal NiV aggregation in hamsters infected with NiV_M_ [34].

**Figure 6.**
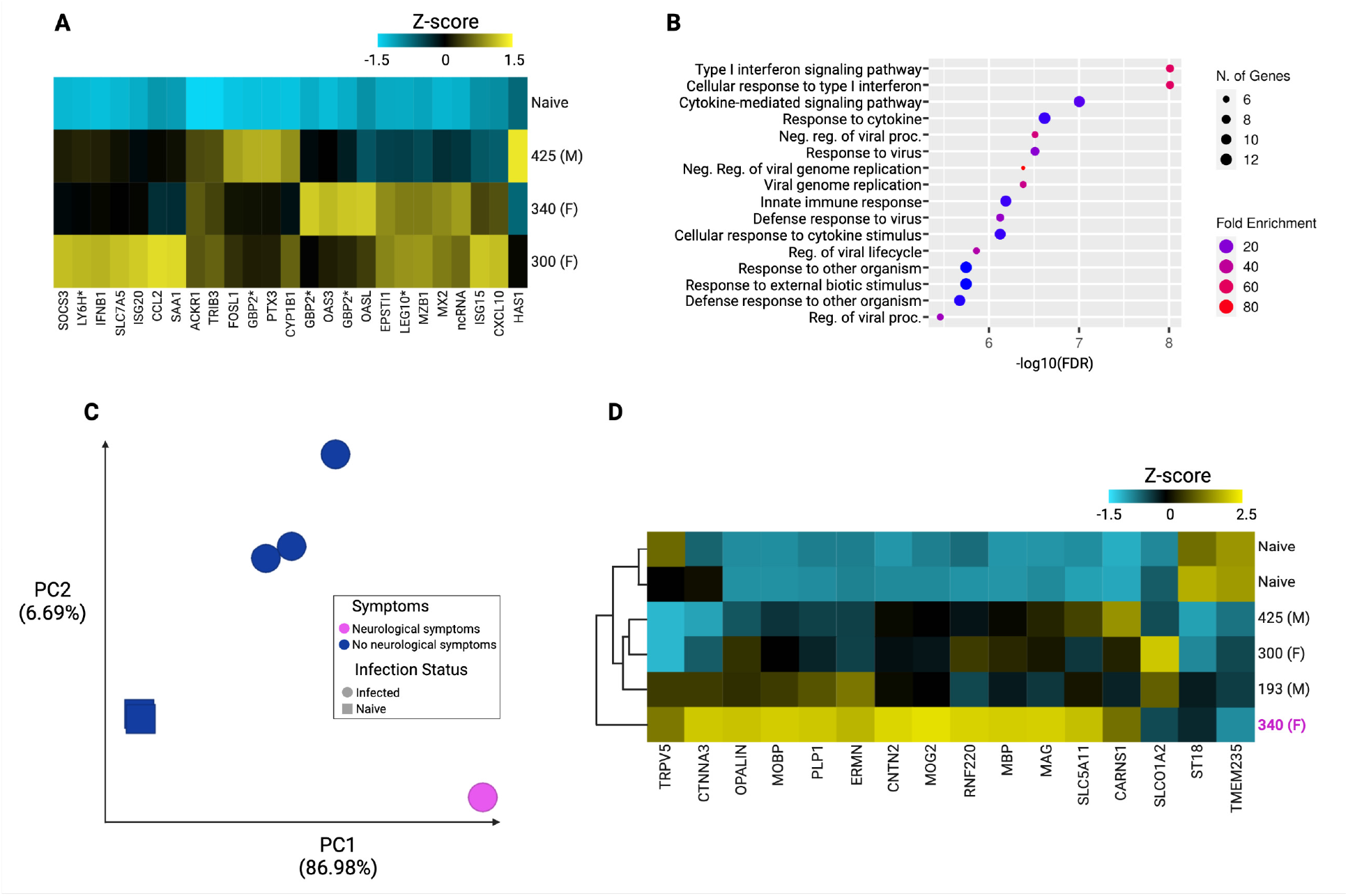
Organ specific responses seen in RNA-seq samples aligned to *Callithrix jacchus*. **(A)** Differential expression analysis across all lung tissue samples was used to identify the top 25 genes most overexpressed in infected versus uninfected lung samples—ranked by FDR step up value—and hierarchical clustering was performed to obtain a heatmap. Gene expression is highest in yellow samples and lowest in cyan samples. **(B)** Enrichment analysis with these 25 genes was performed via ShinyGo 0.76 [41] in order to identify the top 20 most enriched gene sets. **(C)** PCA was performed on all brainstem samples. The top two principal components and their relative contributions are shown. Tissues from infected animals (circles) and tissues from uninfected animals (squares) are further broken down to highlight the brainstem tissue from the only marmoset showing neurological signs (magenta)— marmoset #340 is an outlier by PCA. **(D)** OPALIN, a marker of oligodendrocytes in gene expression profiling [42], and it’s 15 nearest neighbors based on single cell type RNA expression data [43,44] were selected and hierarchical clustering was performed on all brainstem samples. The only marmoset with neurological symptoms (#340, magenta) is a clear outlier and the only marmoset showing high levels of expression of oligodendrocyte-related genes. *An orthologue or “like” protein via the National Center for Biotechnology Information (NCBI).

## Discussion

In this study we identify marmosets as a highly susceptible, novel New World NHP model of NiV_B_ infection. Four marmosets were infected with NiV_B_ via the IN/IT routes and all four animals succumbed to infection by day 11 PI. The marmosets recapitulated many of the same clinical symptoms seen in patients and other NHP models infected with NiV, such as anorexia, lethargy, and hyperventilation. Of special note, one marmoset presented with left hindlimb tremors on day 11 PI, displaying overt neurological clinical symptoms. In the setting of acute infection of NHPs with NiV_B_, neurological clinical symptoms are rare [17] although neurological disease seen via MRI, histopathology, or gross pathology is relatively common [10,14,16,18,20]. Among NHPs neurological clinical symptoms are more common in NiV_M_ than NiV_B_, especially among those surviving longer [9,11,12]. This may give some insight into the development of neurological clinical symptoms in the same marmoset (#340) that presented a less severe infection of the lungs. #340 was found to have a cerebral infarct, malacia, and cerebral edema. This animal, uniquely, also displayed left ventricular cardiomyopathy.

On histopathological examination, all marmosets were found to have significant multifocal coagulation in the liver, syncytia formation of endothelial cells in the kidney, and infiltration in the lung. NiV antigen was found in syncytial cells across multiple tissues, notably in the smooth muscle cells and capillary endothelial cells of the heart. All marmosets also showed consistent hypoalbuminemia and decreases in amylase indicating pancreatic damage. These findings replicate the clinical and pathophysiological picture of NiV infection in NHPs.

Titerable virus was found in all tissue sets evaluated, including the CNS, but some variance was seen across subjects. Similarly, viral RNA was most prevalent in the lower respiratory tract, as expected, but was also found within the CNS for #300, #340, and #425.

RNA-seq provided a deeper insight into both the NiV_B_ and marmoset transcriptional patterns across multiple tissues. NiV_B_ displayed a transcriptional gradient that did not drop until after matrix, a pattern not typically seen among other viruses of the family *Paramyxoviridae* [35,36] but previously noted in vitro with NiV [37]. We also showed that RNA editing of P results in 40% of the total reads being nearly equally split between W and V, in contrast to the often smaller fraction seen in other paramyxoviruses [38,39]. However, our results replicate previous findings specific to NiV_B_ [40]. Similarly, we replicated in vivo findings that the number of G additions is nearly flat from +1 to +5 in HEK293 cells [40].

As expected, across all tissues we observed a consistent up-regulation of gene ontologies relating to regulation of viral processes and interferon stimulated genes. We found results mirroring those found in AGMs with significant upregulation of genes such as *MX2, OASL, CCL2*, and *ISG15* in the most highly infected tissue, the lung [10]. In brainstem samples specifically, the only marmoset with neurologic clinical symptoms was shown to have a unique transcriptional pattern enriched in genes relating to myelination, axon ensheathment, and oligodendrocyte development. Previous findings of myelin degradation and axonal NiV antigen aggregation in hamsters may be related findings to what we observe here in marmosets [34].

Marmosets may provide a new NHP model to study severe disease caused by NiV. These findings are notable given many of the logistical advantages afforded by working with marmosets in the context of high-biocontainment human respiratory pathogens. Marmosets offer distinct advantages to the current standard NHP model of NiV infection, AGMs, due to their small size and ease of handling, typically lower costs, and ease of co-housing allowing for a potential model of transmission studies. Given the unique transcriptional profile in the brainstem of the only marmoset that showed neurologic clinical symptoms, there is a need for additional follow up with a larger cohort number of both infected and uninfected marmosets. More detailed pathogenicity studies are required to monitor levels of viremia, as well as evaluate different routes of infection to potentially allow in depth studying of neuroinvasion. Overall, marmosets were highly susceptible to infection via the IN/IT route resulting in a 100% mortality in non-protected animals. (3435 words)

## Acknowledgments

We thank the University of Texas Medical Branch Animal Resource Center for husbandry support of laboratory animals. Figures created with BioRender.com.

## Notes

### Funding statement

This work was supported by the National Institutes of Health [U19 AI171403 to B.L. and A.F] and the National Institutes of Health Viral-Host Pathogenesis Training Grant [T32 AI07647 to C.S.].

Portions of this data have been previously presented at the 2019 Nipah Virus International Conference (Singapore), the American Society for Virology Meeting 2021 (virtual), and the Negative Strand Virus Conference 2022 (Braga, Portugal).

