## Supplemental Figures for "Nipah virus Bangladesh infection elicits organ-specific innate and inflammatory responses in the marmoset model"

| Subject No. | Sex | Euthanized (DPI) | Clinical and gross pathology | Clinical diagnosis |
|---|---|---|---|---|
| **193** | Male | 11 | (1) Minimal (<10%) normal lung tissue detected in the cranial portion of the lung. The rest of lung was fluid filled with hemorrhagic/serous fluid. The left and right bronchus had clear fluid present and the trachea had minimal fluid in it.<br>(2) Liver had multi-focal to coalescing regions of pale circles of varying sizes throughout all fields and was very friable.<br>(3) Mediastinal and bronchial lymph nodes had gross enlargement and hemorrhage. | (1) Diffuse Pulmonary Edema and Hemorrhage<br>(2) Diffuse hemorrhagic hepatitis<br>(3) Multi-focal hemorrhagic Lymphadenopathy |
| **300** | Female | 9 | (1) Minimal (<5%) normal lung tissue detected in the cranial portion of the lung. The rest of lung was fluid filled with hemorrhagic/serous fluid. The left and right bronchus had serosanguineous fluid present, and the trachea had minimal fluid in it.<br>(2) Liver had multi-focal to coalescing regions of pale circles of varying sizes throughout all fields and was very friable.<br>(3) Inguinal and retropharygneal lymph nodes had gross enlargement but minimal hemorrhage associated with them.<br>(4) Tonsils had pinpoint hemorrhages associated throughout the tissue. The tissue however was not enlarged.<br>(5) Diffuse subcutaneous liquefactive necrosis of fat.<br>(6) Thoracic cavity contained approximately 1mL of serosanguineous fluid.<br>(7) Peritoneal cavity contained approximately 2mL of serosanguineous fluid. | (1) Diffuse Pulmonary Edema and Hemorrhage<br>(2) Diffuse hemorrhagic hepatitis<br>(3) Multi-focal Lymphadenopathy<br>(4) Hemorrhagic Tonsilitis |
| **340** | Female | 11 | (1) Right lung was normal, though the bronchus was fluid filled. Left lung was congested as it was the dependent lung at the time of death and is a post-mortem change.<br>(2) Liver had multi-focal to coalescing regions of pale circles of varying sizes throughout all fields and was very friable.<br>(3) Inguinal lymph node had gross enlargement, but minimal hemorrhage associated with it.<br>(4) Significant hypertrophy of the left ventricular (LV) wall, approximately 10:1 left vs. right. Dilation of the right atrium, though milder than the LV hypertrophy.<br>(5) Infarct was found and malacia near the transition zone between grey and white matter in the occipital lobe of the left cerebral hemisphere. Edema was appreciated in the cerebrum.<br>(6) Trachea was filled with serous fluid at necropsy.<br>(7) Multiple pinpoint areas of hemorrhage were appreciated in the right renal pelvis.<br>(8) Peritoneal cavity contained approximately 0.5 mL of serosanguineous fluid. | (1) Diffuse hemorrhagic hepatitis<br>(2) Left Ventricular Cardiomyopathy / Right Atrial Dilation<br>(3) Cerebral Infarct and malacia in the occipital lobe<br>(4) Cerebral Edema |
| **425** | Male | 8 | (1) Minimal (<5%) normal lung tissue detected in the cranial portion of the lung. The rest of the lung was fluid filled with hemorrhagic/serous fluid. The left and right bronchus had clear fluid present and the trachea had minimal fluid in it.<br>(2) Liver had multi-focal to coalescing regions of pale circles of varying sizes throughout all fields and was very friable.<br>(3) Inguinal, mediastinal and retropharygneal lymph nodes had gross enlargement and hemorrhage.<br>(4) Spleen was extremely friable and congested, no normal tissue was observed on cut sections. | (1) Diffuse Pulmonary Edema and Hemorrhage<br>(2) Diffuse hemorrhagic hepatitis<br>(3) Multi-focal hemorrhagic Lymphadenopathy<br>(4) Splenic Congestion |

**Table S1. Clinical description and diagnosis in $NiV_B$ infected marmosets.**

Detailed descriptions of clinical and gross pathology as well as clinical diagnoses for all four marmosets infected with $NiV_B$.

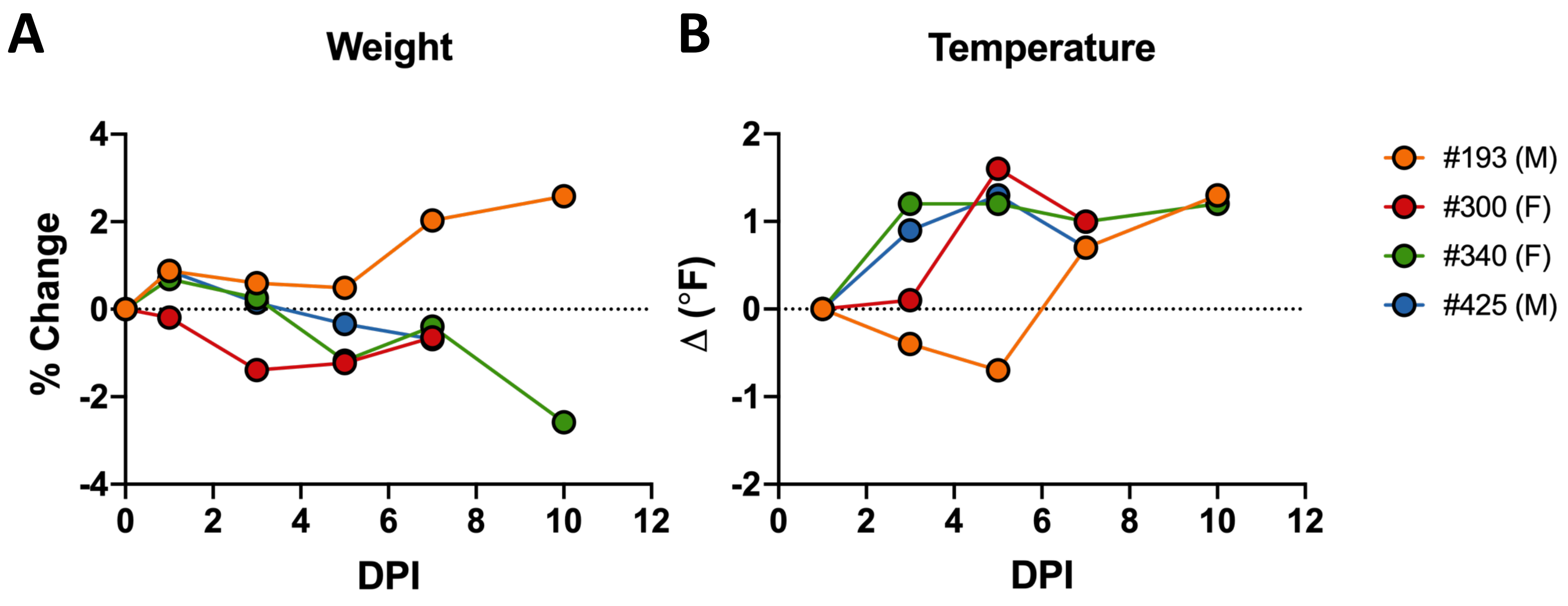


**Figure S1: Changes in body weight and temperature in marmosets after $NiV_B$ infection.**
(A) Changes in body weight are shown as percent change compared to the individual starting weight at day 0 of the study. (B) Changes in body temperature are shown as changes in ºF compared to the first measurement made on day 1 post-infection. Subject #193 is shown in orange circles, subject #300 in red circles, subject #340 in green circles, and subject #425 in blue circles.

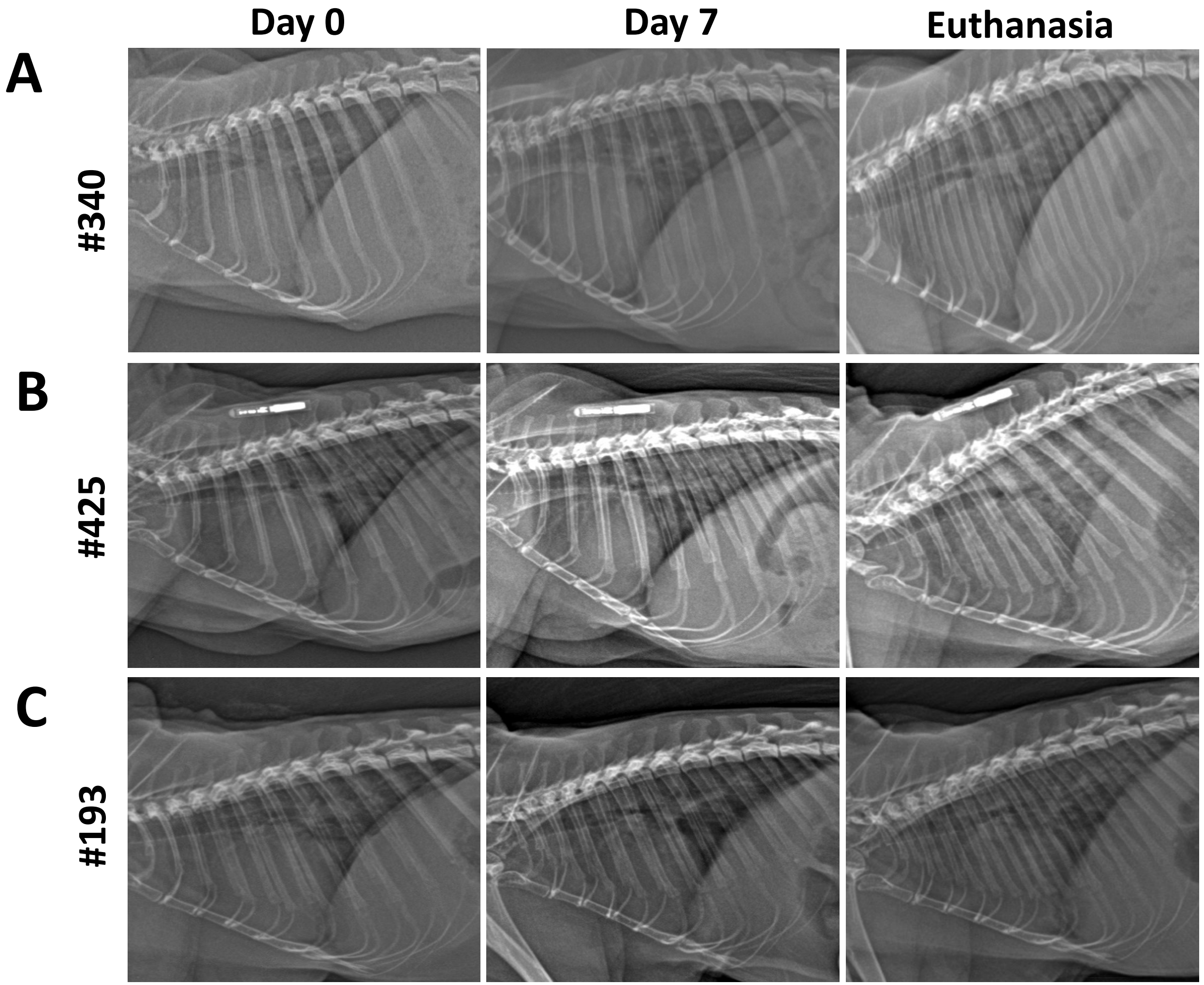


**Figure S2: Progressive lung pathogenesis in marmosets after $NiV_B$ infection.**

Subject were positioned in a left lateral position and all images are in a Cranial to Caudal orientation. X-ray images were taken on days 0, 1, 3, 5, 7, and at euthanasia. Prior to day 7, lungs appeared normal. Progressive lung pathology was detected from day 7 onward. **(A)** Subject #340: A mild bronchial pattern emerged at the time of euthanasia (day 9 post-infection) and showed some inflammation around the smaller bronchi, compared to day 7. **(B)** Subject #425: A severe interstitial pattern with nodules emerged at the time of euthanasia (day 11 post-infection), compared to day 7 where the animal was showing some inflammation around the smaller bronchi with a mild interstitial infiltrate. **(C)** Subject #193: A severe interstitial pattern emerged at the time of euthanasia (day 11 post-infection), compared to day 7 where the lung was completely normal.

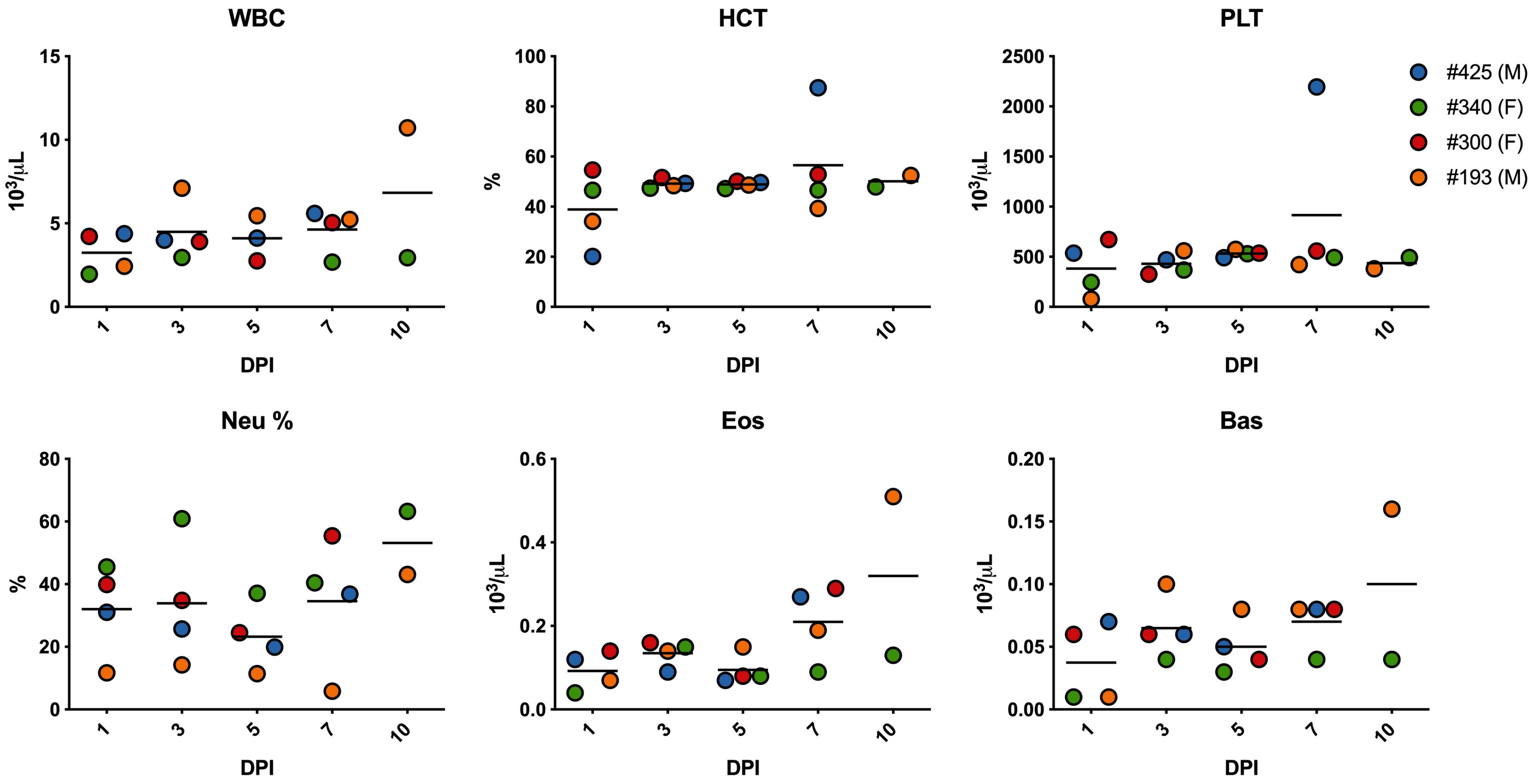


**Supplementary Figure S3. Changes in complete blood count.**

Taken at days 1, 3, 5, 7, and 10, hematological analysis included total white blood cell counts, white blood cell differential, red blood cell counts, platelet counts, hematocrit values, total hemoglobin concentrations, mean cell volumes, mean corpuscular volumes and mean corpuscular hemoglobin concentrations. White blood cell count (WBC), neutrophils (Neu), eosinophils (Eos), basophils (Bas), hematocrit (HCT), and platelets (PLT).

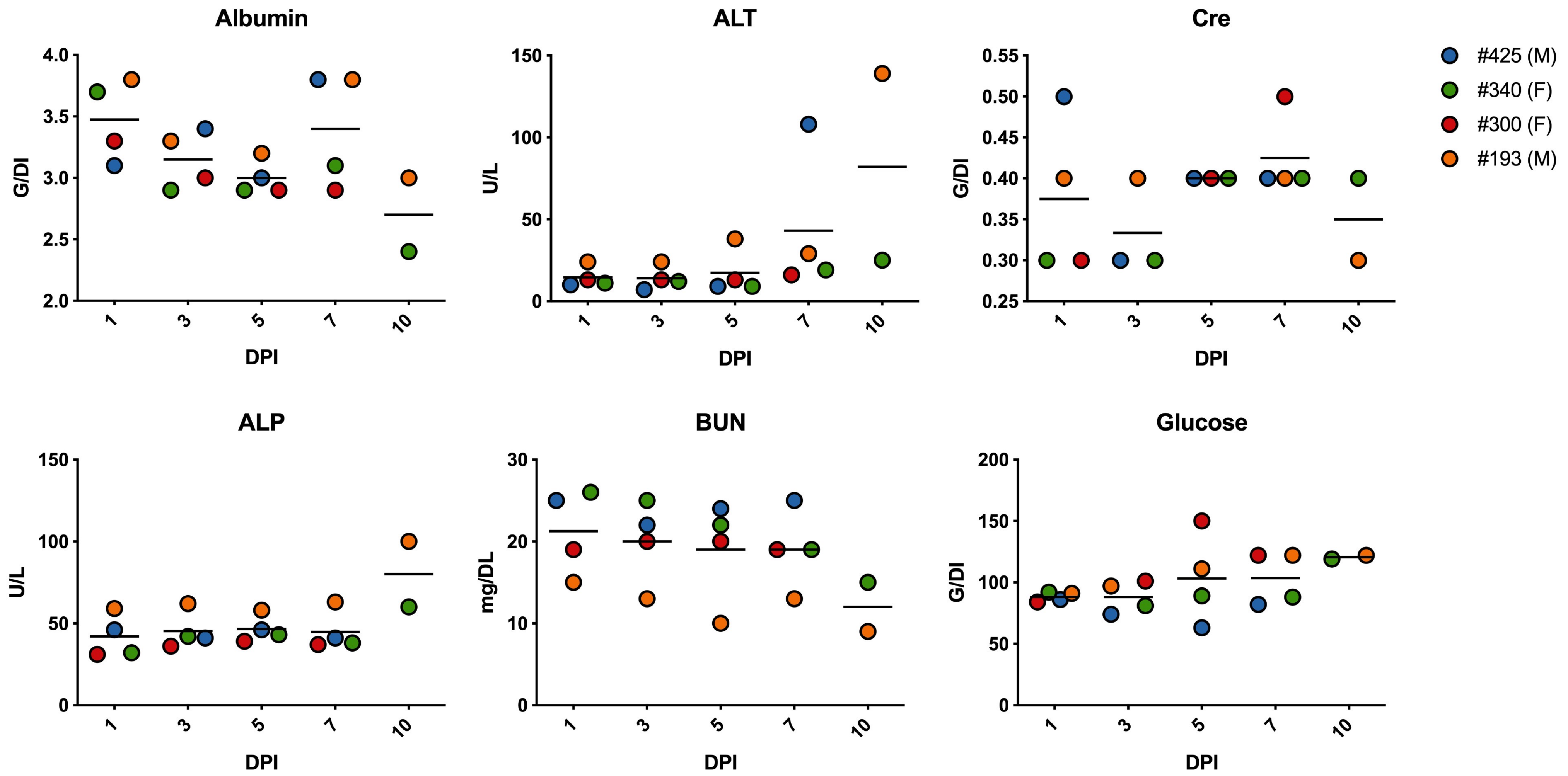


**Supplementary Figure S4. Changes in blood chemistry.**

Clinical chemistry analysis was performed at days 1, 3, 5, 7, and 10. Serum samples were tested for concentrations of albumin, alanine aminotransferase (ALT), alkaline phosphatase (ALP), glucose, plasma electrolytes (blood urea nitrogen (BUN) and creatinine (CRE)).

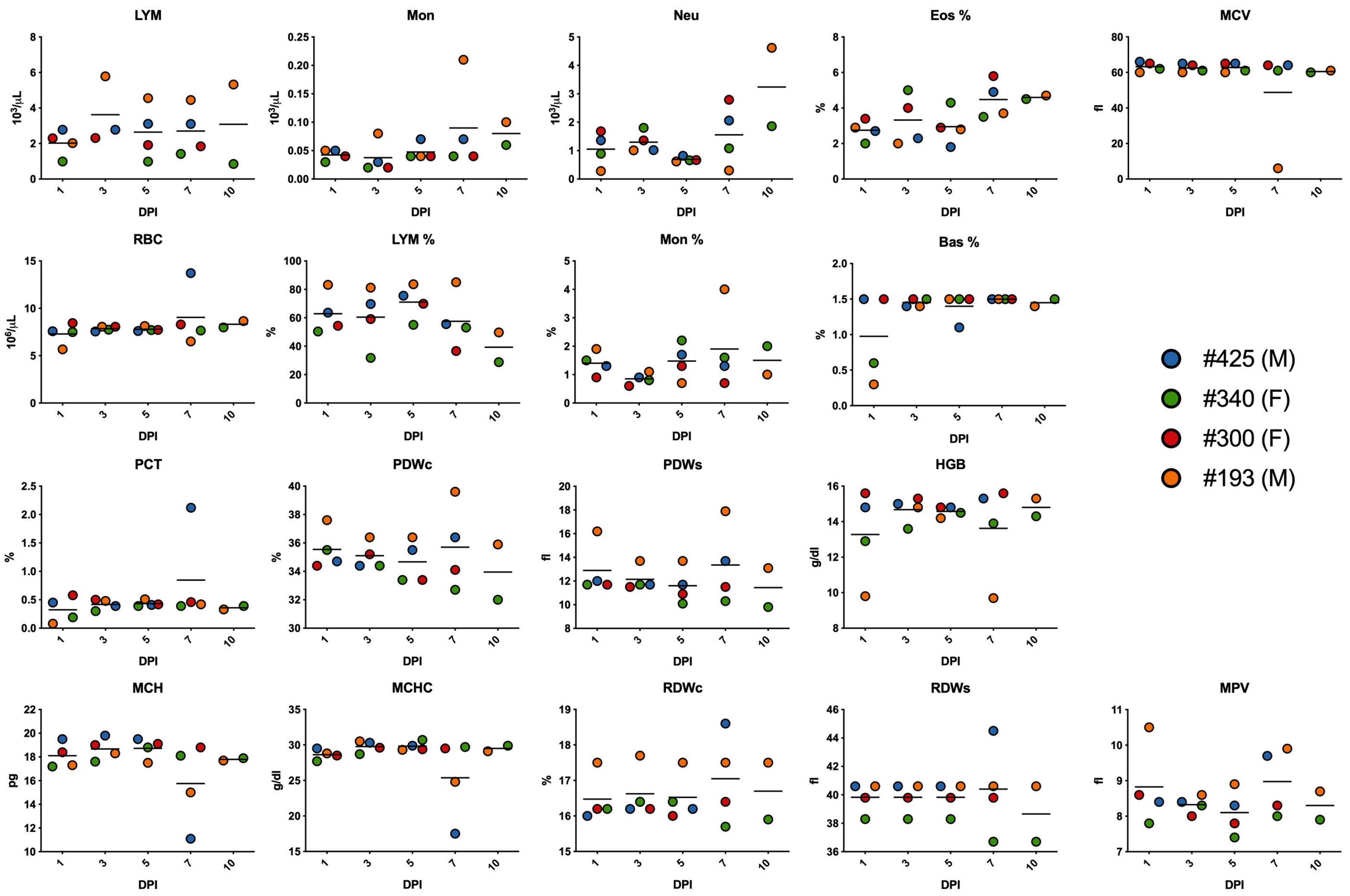


**Supplementary Figure S5. Additional changes in complete blood count.**

Taken at days 1, 3, 5, 7, and 10, hematological analysis included total white blood cell counts, white blood cell differential, red blood cell counts, platelet counts, hematocrit values, total hemoglobin concentrations, mean cell volumes, mean corpuscular volumes and mean corpuscular hemoglobin concentrations. Red blood cell count (RBC), lymphocytes (LYM), monocytes (Mon), plateletcrit (PCT), platelet distribution width (PDWs), platelet distribution width % (PDWc), hemoglobin (HGB), mean corpuscular volume (MCV), mean corpuscular hemoglobin (MCH), mean corpuscular hemoglobin concentration (MCHC), red cell distribution width (RDWs), red cell distribution width % (RDWc), and mean platelet volume (MPV).

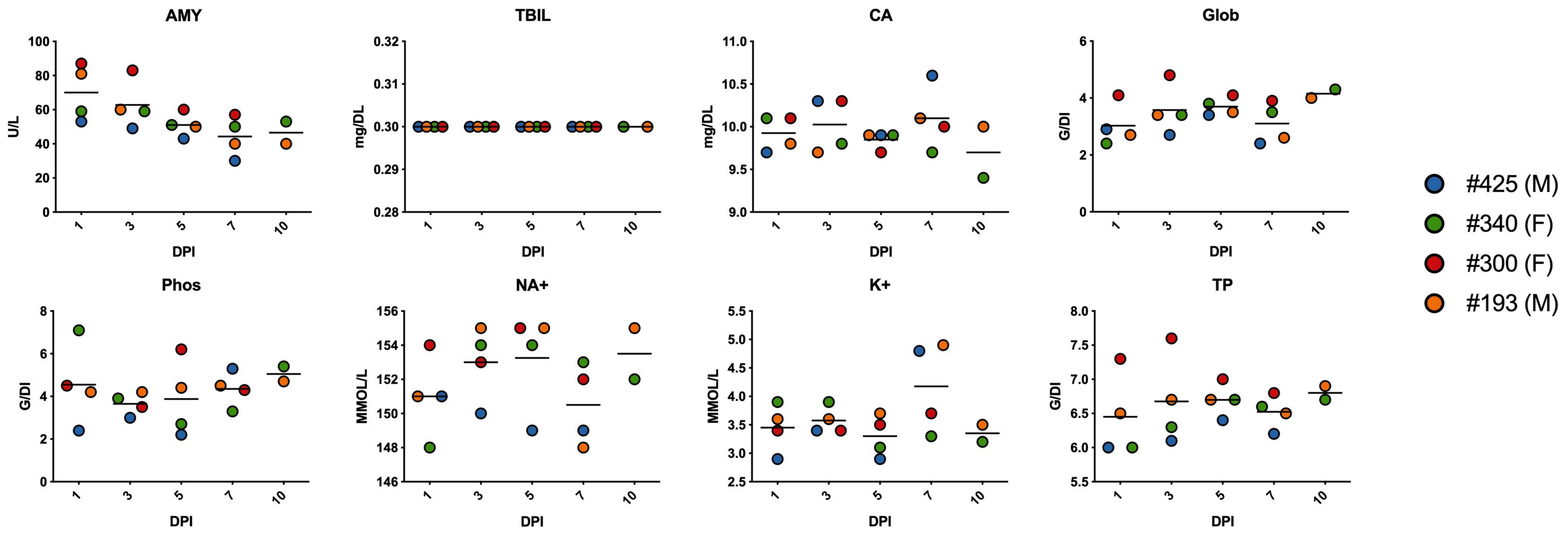


**Supplementary Figure S6. Additional changes in blood chemistry.**

Clinical chemistry analysis was performed at days 1, 3, 5, 7, and 10. Serum samples were tested for concentrations of amylase (AMY), total protein (TP), total bilirubin (TBIL), plasma electrolytes (calcium (CA), phosphorus (Phos), sodium (Na+), and potassium (K+), and globulin (GLOB)).

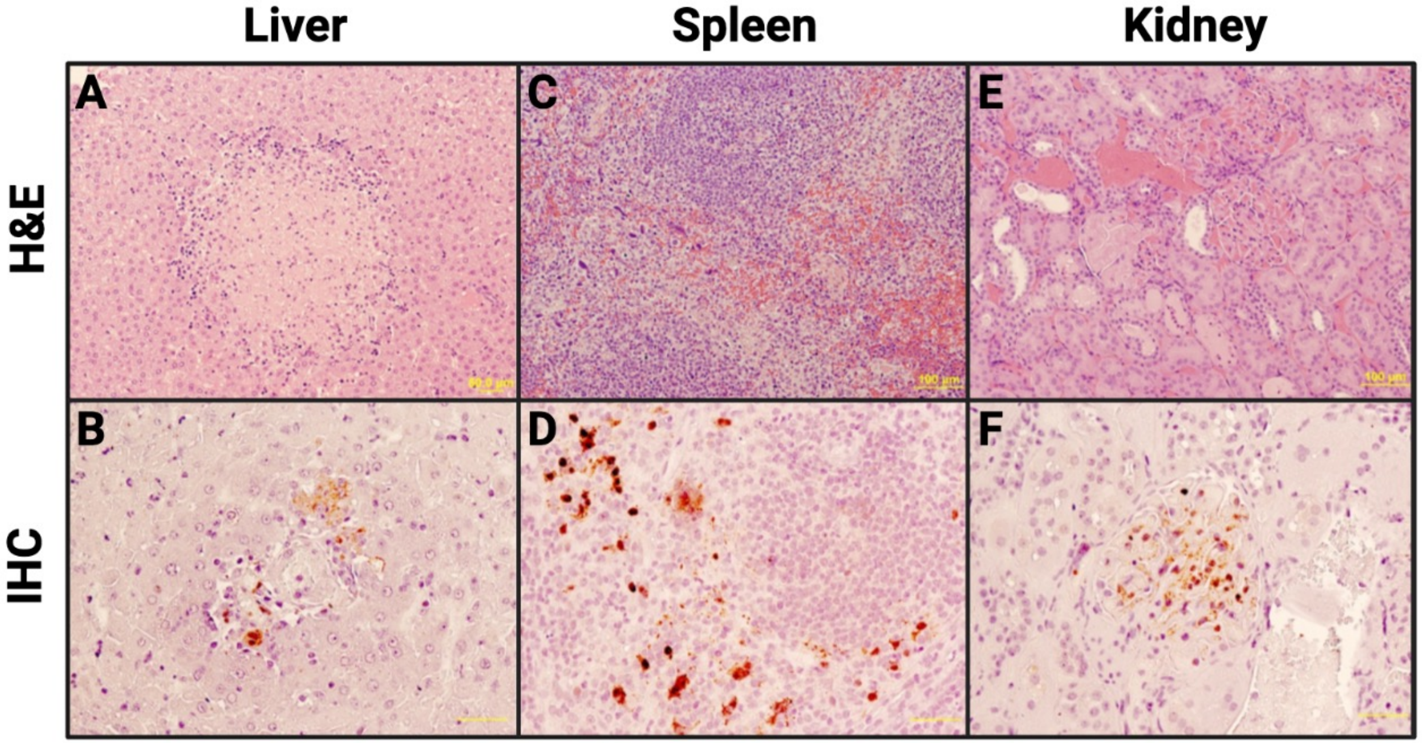


**Supplementary Figure S7: Additional histopathological changes in marmoset tissues after NiV$_B$ infection.**
**Panels A, C, E:** H&E staining**. Panels B, D, F:** Immunohistochemical staining for NiV nucleoprotein.
(**A**) Liver (subject 340): Focal coagulation necrosis in midzonal hepatocytes, surrounded by inflammatory cells (neutrophils, mononuclear cells). (**B**) Liver (subject #425): NiV antigen present in hepatocytes, sinusoidal endothelial cells, and portal vein endothelial cells. **(C)** Spleen (subject #340): Necrosis of red pulp and formation of syncytial cells. **(D)** Spleen (subject #425): NiV antigen present in mononuclear or multinuclear cells in peripheral zone of follicles, or in red pulp. **(E)** Kidney (subject #300): Loss of endothelial and mesangial cells in affected glomeruli. **(F)** Kidney (subject #425): NiV antigen present in glomerular cells and peritubular capillary endothelial cells.
